# *In silico* hippocampal modeling for multi-target pharmacotherapy in schizophrenia

**DOI:** 10.1101/758466

**Authors:** Mohamed A Sherif, Samuel A Neymotin, William W Lytton

## Abstract

**Background:** Treatment of schizophrenia has had limited success in treating core cognitive symptoms. The evidence of multi-gene involvement suggests that multi-target therapy may be needed. Meanwhile, the complexity of schizophrenia pathophysiology and psychopathology, coupled with the species-specificity of much of the symptomatology, places limits on analysis via animal models, *in vitro* assays, and patient assessment. Multiscale computer modeling complements these traditional modes of study.

**Methods:** Using a hippocampal CA3 computer model with 1200 neurons, we examined the effects of alterations in NMDAR, HCN (*I*_h_ current), and GABA_A_R on information flow (measured with normalized transfer entropy), and in gamma activity in local field potential (LFP).

**Results:** Altering NMDARs, GABA_A_R, *I*_h_, individually or in combination, modified information flow in an inverted-U shape manner, with information flow reduced at low and high levels of these parameters. The strong information flow seen at the peaks were associated with an intermediate level of synchrony, seen as an intermediate level of gamma activity in the LFP, and an intermediate level of pyramidal cell excitability.

**Conclusions:** Our results are consistent with the idea that overly low or high gamma power is associated with pathological information flow and information processing. These data suggest the need for careful titration of schizophrenia pharmacotherapy to avoid extremes that alter information flow in different ways. These results also identify gamma power as a potential biomarker for monitoring pathology and multi-target pharmacotherapy.

**AUTHOR SUMMARY:** Currently, there are no good treatments for the cognitive symptoms of schizophrenia. We used a biophysically realistic computational model of hippocampal CA3 to investigate the effect of potential pharmacotherapeutic targets on the dynamics of CA3 activity and information processing to predict multi-target drug treatments for schizophrenia. We found an inverted-U shaped relationship between information flow and drug target manipulations, as well as between information flow and gamma power. Our study suggests that neuronal excitability and synchrony may be tuned between extremes to enhance information flow and information processing. It further predicts the need for careful titration of schizophrenia drugs, whether used individually or in drug cocktails.

## 1 Introduction

Schizophrenia is a chronic disease with a lifetime prevalence of around 4/1000 [75], which usually produces life-long disability [38]. Cognitive impairment and information processing deficits are chief causes of disability [56, 36, 22]. The most affected cognitive domains are processing speed, working memory, episodic memory, and verbal learning and memory [18, 6, 24]. Current antipsychotic medications have limited impact on cognitive symptoms and information processing deficits [8, 14]. Therefore, there are significant gaps in therapy and patients’ clinical care [20].

Recent research has emphasized the role of glutamatergic transmission as an extension of the dopaminergic hypothesis for schizophrenia pathophysiology, especially to capture cognitive impairment associated with schizophrenia (CIAS) and information processing deficits [57]. The role of glutamatergic transmission has been supported by the psychotomimetic effect of N-methyl-D-aspartate (NMDA) receptor (NMDAR) antagonists like phencyclidine (PCP) and ketamine in healthy volunteers [31, 44]. Ketamine also worsened cognitive symptoms in schizophrenia patients [55].

Gamma oscillations have been implicated in a number of brain functions, including sensory integration [93]. Gamma is hypothesized to enable local computations within cortical microcircuits [3] by providing binding among neurons belonging to a particular neuronal ensemble, which would fire synchronously with this gamma rhythm [16, 52]. These oscillations may also play a role in routing information across brain regions [15]. Patients diagnosed with schizophrenia have shown abnormalities in gamma oscillations, with multiple studies showing a reduction in induced and evoked gamma power [35, 42, 50, 87]. By contrast, pre-symptomatic, clinically high-risk individuals demonstrated an increase in resting gamma power [23]. Ketamine increased resting gamma power in healthy volunteers [71]. Interestingly, acute ketamine increased resting gamma power but reduced induced gamma in rodent hippocampal CA3 [45], while chronic ketamine reduced gamma power [37], similar to what has been reported in patients with chronic schizophrenia [23].

Combining evidence from gamma studies and ketamine studies suggests a role for NMDAR dysfunction in gamma abnormalities and in CIAS. Additional evidence comes from the schizophrenia genome-wide association study (GWAS) published in 2014, which identified GRIN2A (glutamate ionotropic NMDA-type receptor subunit 2A) on chromosome location 16p13 as being close to one of the 108 gene loci identified [78]. Clinically, agonists for NMDARs that have been studied for the treatment of CIAS include glycine and D-serine, both of which bind to an allosteric site on the GluN1 subunit of NMDAR and are obligatory for NMDAR activation by glutamate [58]. D-cyclo-serine, a partial agonist at the glycine site [79], has also shown efficacy in some studies. However, these studies yielded mixed results; a recent meta-analysis found no overall significant effect on cognition, although young patients aged 30 to 39 years old showed some benefit [11].

Another molecular aspect that we consider in this paper is the involvement of the HCN1 gene (hyperpolarization-activated cyclic nucleotide-gated channel type 1) that codes for *I*_h_ channels (hyperpolarization-activated current, also known as anomalous rectifier, *I*_f_, *I*_q_) [1, 12, 77]. Along with NMDAR, this channel plays a role in the generation and modulation of neuronal oscillations [60, 62]. The HCN1 gene on chromosome location 5p21 is also close to one of the GWAS loci associated with schizophrenia [78]. The role of *I*_h_ in oscillations, and evidence from the GWAS study, suggest HCN1 product as another potential therapeutic target for CIAS [34].

Manipulating the GABA_A_ system provides another potential treatment target [94]. The GABAergic system shapes synchronized neuronal activity during oscillations [54, 13, 80, 69, 73], and is impaired in schizophrenia. Post-mortem studies have revealed low inhibitory interneuron glutamic acid decarboxylase enzyme (GAD67), low GABA transporter, and low pyramidal cell GABA_*A*_ receptor (GABA_A_R) *α*1 subunit mRNA transcripts in the frontal lobe of patients [25]. Similar findings were also demonstrated in the hippocampus, where a reduction in numbers of somatostatin-positive and parvalbumin-positive interneurons was also found [41]. The GABAergic deficit hypothesis for schizophrenia is further supported by reduced post-mortem immunoreactivity of GAD65/67 in interneuronal neuropil in the hippocampus [82]. Reduced GABA tone has been suggested to mediate hippocampal hyperactivity in these patients [26, 49, 84, 85, 86].

Schizophrenia pathophysiology is extremely complex, with abnormalities at multiple scales, from genes, second messenger cascades, and cells, up to local networks and inter-areal communication. Given this complexity, it is reasonable to consider that multi-target pharmacotherapeutic approaches will be useful [40]. Targets in multi-target pharmacotherapy interact in highly non-linear ways, making it impossible to intuitively predict the effects [59, 53, 63]. We, therefore, use simulations to study these interactions. In this study, we investigated how alterations in the three targets mentioned above – NMDAR, *I*_h_, GABA_A_R – will affect 1. oscillations and 2. information flow in a biophysically-realistic computer model of the CA3 region of the hippocampus. This builds on our prior results which identified the effects of blocking NMDARs on oriens-lacunosum moleculare (OLM) interneurons on gamma oscillations and information flow [66].

## 2 Results

### 2.1 Baseline network activity produced theta with nested gamma oscillations

Around 1900 simulations were run using NEURON 7.4 [10, 28]. Individual simulations, run for seven seconds model time, required about five minutes, running in parallel on a Linux system with eight 2.67 GHz Intel Xeon quad-core CPUs. We investigated the correlations between ion channel manipulations, scaling 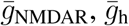, and 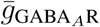, with information flow and gamma oscillation strength. Multiple versions of individual simulations with different randomizations were run to ensure that results were generalizable, and not due to specific network architecture or network input.

In baseline simulations (Fig. 1), firing of the three neuronal populations was synchronized at two primary frequencies: gamma (∼25 to 50 Hz) and theta (∼6 to 8 Hz). Synchronized activity resulted from the following sequence of events: pyramidal neuronal firing triggered firing of PV basket interneurons, which in turn turned off the pyramidal neurons until inhibition wore off (pyramidal interneuron network gamma, PING oscillations). Pyramidal neurons also triggered the firing of OLM interneurons, which provided further feedback inhibition to the pyramidal neurons, but at a slower rhythm (theta) due to OLM interneurons’ longer-lasting inhibition from longer GABA_A_R time constants. Synchronization at the theta frequency was also mediated by inhibitory input from the medial septum to both OLM and PV interneurons, but was not dependent on it. While pyramidal neurons were under the inhibitory influence of the OLM interneurons, the firing and reciprocal inhibition between PV basket interneurons generated interneuron network gamma (ING) oscillations. The synchronized firing of the pyramidal neurons, PV basket, and OLM interneurons was reflected in the local field potential (LFP) (Fig. 1B). The LFP showed both gamma and theta frequencies. The power spectral density (PSD) (Fig. 1C) showed the peaks of gamma (∼ 35 Hz) and theta (∼ 7 Hz) oscillations in this representative simulation.

**Figure 1:**
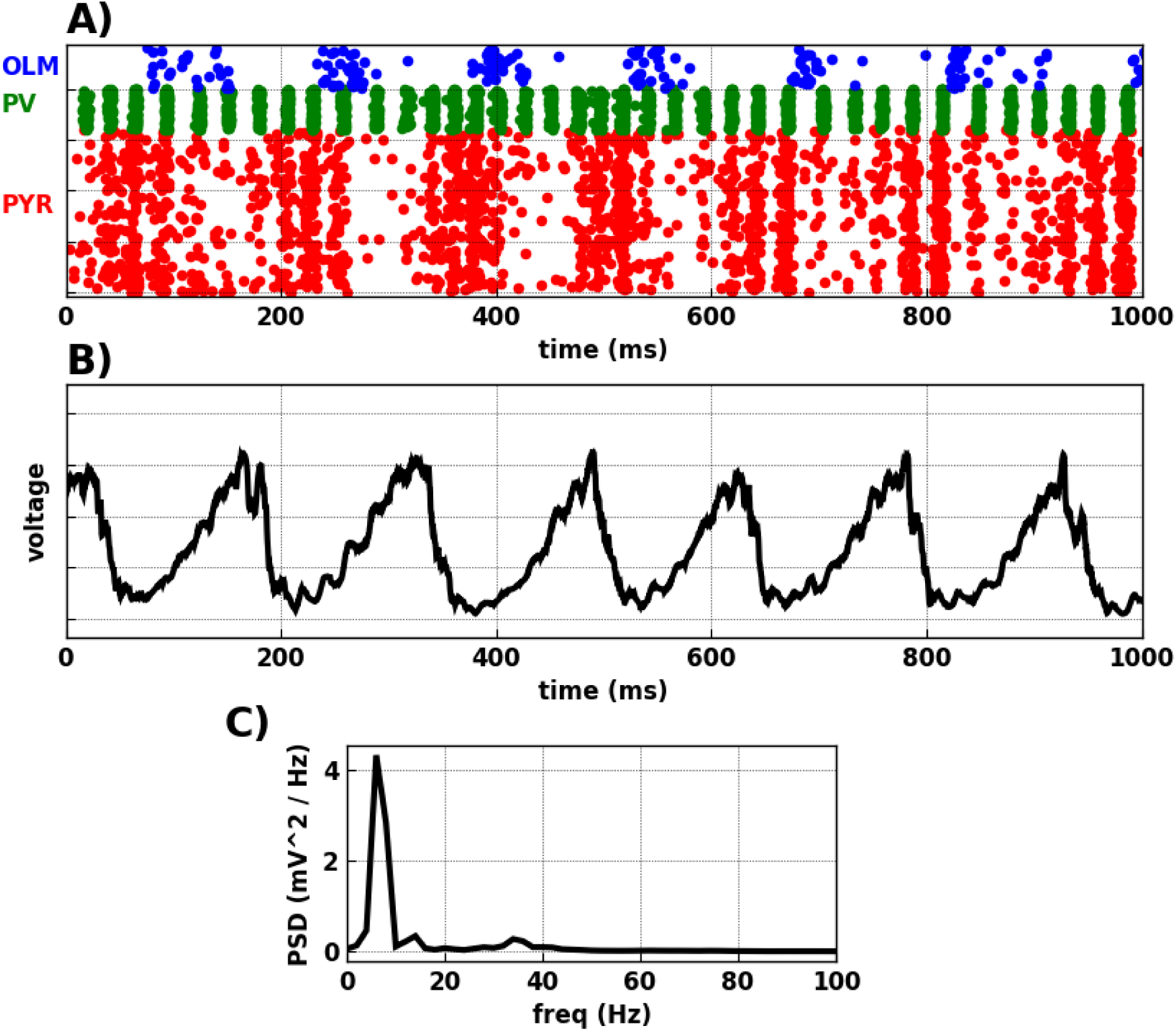
Baseline simulation generated theta (6-8 Hz) with nested gamma (30-40 Hz) rhythm. **(A)** Raster plot where each row represents a neuron and each dot represents an action potential (x-axis: time; y-axis: neuron identity; OLM interneurons in blue, PV basket interneurons in green, pyramidal neurons in red). **(B)**. Simulated local field potential (LFP). **(C)** Power spectral density (PSD) of LFP showed theta (∼ 7 Hz) and gamma (30-40 Hz) peaks.

### 2.2 Augmenting OLM NMDARs modulated circuit oscillations and information flow

Our first step was studying the effects of augmenting NMDAR of OLM interneurons on gamma oscillations and information flow across the model (Fig. 2). OLM interneurons were the location where NMDAR antagonism produced theta and gamma oscillatory changes similar to what was seen with ketamine in hippocampal CA3 [66, 45]. We scaled 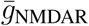 of OLM interneurons from 1.5 *×* to 30 *×* the control simulations in Fig. 1. This scaling could reflect a conductance increase due to phosphorylation or insertion of different isoforms with or without an increase in the number of NMDAR channels. Scaling 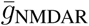 of OLM interneurons reduced gamma power ∼10-fold until it disappeared at above 30*×* control (Fig. 2B, left y-axis).

**Figure 2:**
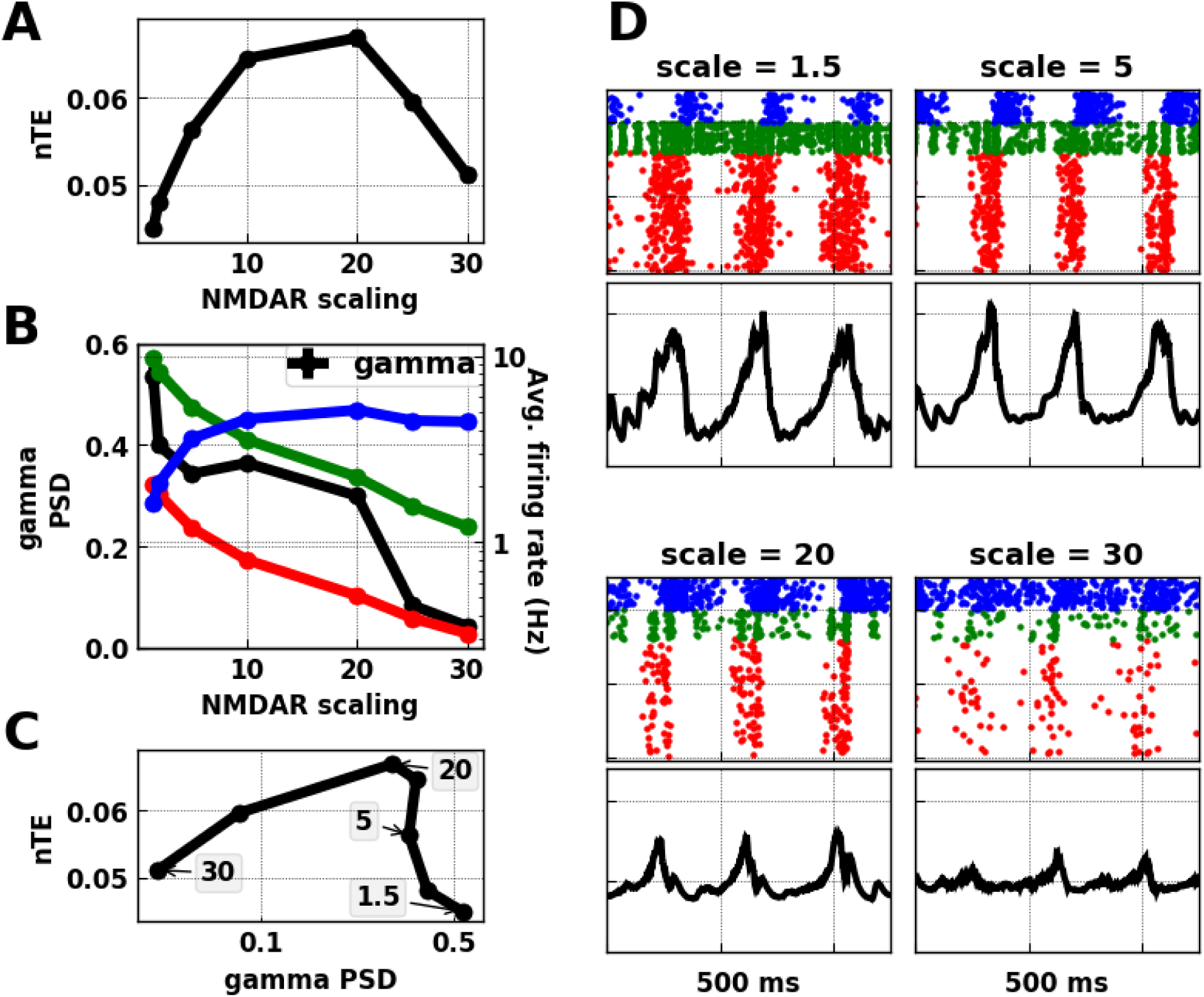
Augmenting 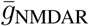 on OLM interneurons. A. Information transfer (nTE) shows an inverted-U relationship with scaling OLM 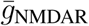 up (mean *±* SEM). **B.** Increasing OLM 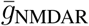 increased activity of OLM interneurons (blue), reduced pyramidal (red) and PV (green) neuronal firing, and reduced gamma power (mean *±* SEM). **C.** Inverted-U relationship between gamma power and information flow shown with different levels of OLM 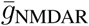 (mean *±* SEM). Note that highest 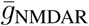 is now at the left (values in shaded rectangles). **D.** Raster plots and LFP at different 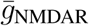 scaling relative to control.

Scaling OLM 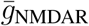 upwards increased drive onto OLM interneurons and therefore increased the firing rate of these inhibitory cells (Fig. 2B, blue line), which reduced the firing of the other two cell types that received OLM projections (PV, green; PYR, red). Information transfer (nTE) across the excitatory (PYR) population (Fig. 2A) peaked at an intermediate level of PYR and PV firing, and showed far lower values with firing rate extremes, low or high. We found a similar inverted-U that related information flow to gamma power, a measure of synchronized firing (Fig. 2C). Note that the values of 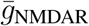 are reversed in **C** relative to **A** and **B**, with high 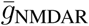 at left associated with low PYR firing and low gamma. There were two causes for low gamma with low PYR rate: **1.** reduced number of spikes, which also reduced the strength of theta; **2.** reduction in spiking coherence which was no longer as sharply shaped by PING and ING (Fig. 2D, scale=30).

### 2.3 Modulating information flow with multiple channel manipulations

Consideration of multi-target pharmacotherapy requires determining how actions at different targets combine. We, therefore, looked at alterations in *I*_h_ and GABA_A_R, both of which are considered possible factors in schizophrenia pathophysiology [41, 78], along with 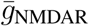. Having previously demonstrated strong effects on gamma of HCN channels of pyramidal and PV neurons [60], we focused here on *I*_h_ at these two locations. The inverted-U configuration of the nTE peak for 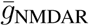 augmentation was seen at all *I*_h_ scalings except for 10 *×* (Fig. 3). An inverted-U pattern could also be seen for GABA_A_R scaling with 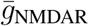 10 *×*, again at all but the highest *I*_h_ value. Similarly, the inverted-U peak can be detected for *I*_h_ scaling around the point marked H1, with a less well-defined, broader surface peaking around the point marked by H2. Thus, we have identified parameter ranges that identify a set of peaks which are points of strong nTE in a 4-D space based on this 3-D parameter space.

**Figure 3:**
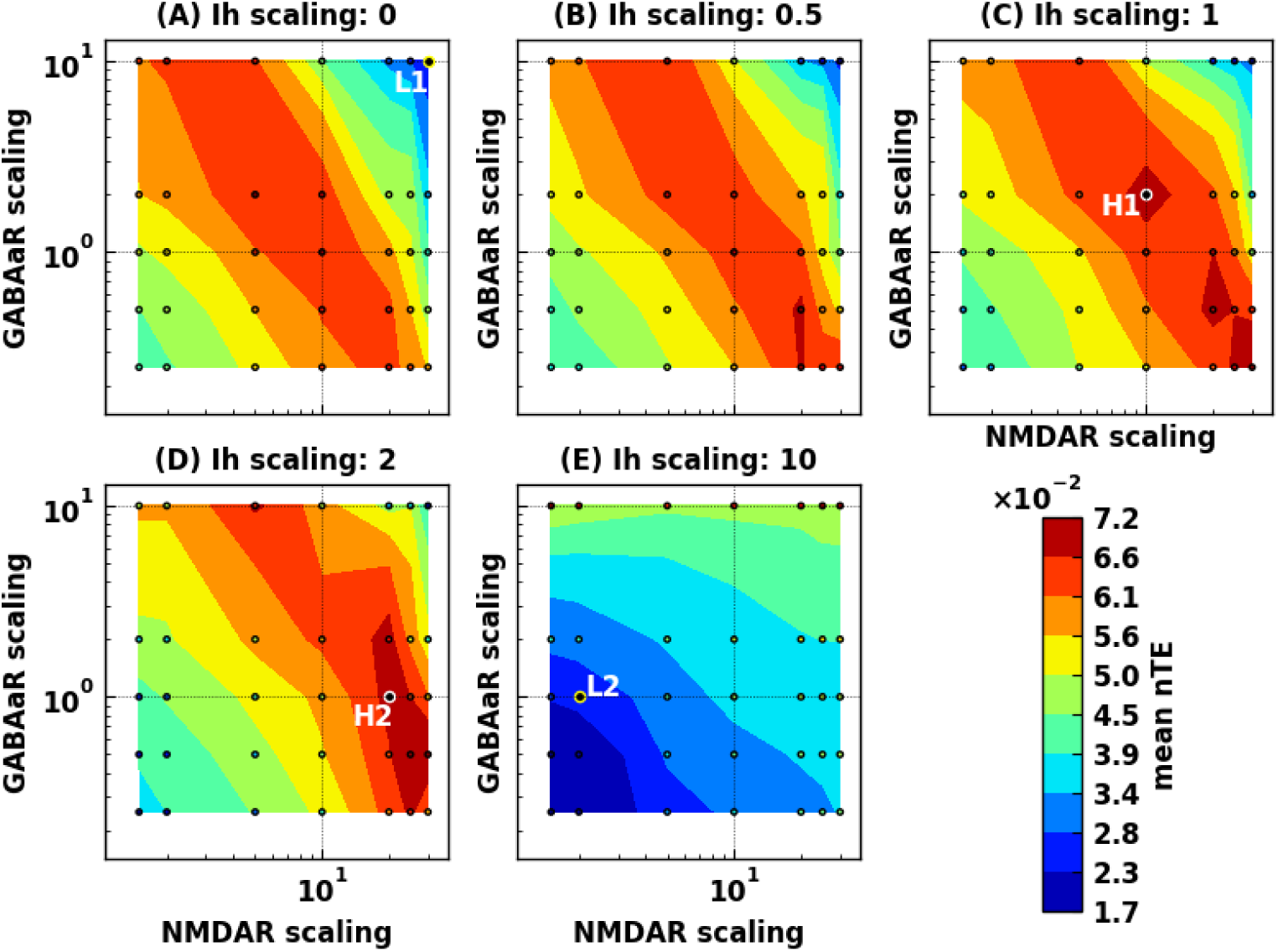
Multi-target manipulations modulate information flow. *I*_h_ scaled in panels **A** - **E**, with 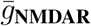 and 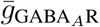 on x- and y-axis in each case. Values are interpolated from simulation results (small circles).

Firing patterns from regions of low (L1-2; Fig. 4) and high (H1-2; Fig. 4) nTE were easily distinguishable. Two patterns of neuronal firing were seen which produced low nTE: L1. low firing; L2. over-synchrony. L1 exhibited low oscillatory power in addition to low firing. This was because of the combined inhibitory effect of high NMDAR scaling on OLM interneurons (30 *×*) and high GABAaR scaling on pyramidal neurons (10 *×*), leading to inhibition of pyramidal neurons and PV basket interneurons. In L2, highly-synchronized phase-locked pyramidal neuron firing was driven by strong PV basket feedback inhibition, permitting little of the variation required to transfer information. This highly-synchronized dynamics resulted from high *I*_h_ scaling (10 *×*) on pyramidal neurons and PV interneurons, increasing their excitability and their synchronous activity, similar to what we reported before [64]. The different dynamics between L1 and L2 were reflected in their gamma power (bottom panel of Fig. 4).

**Figure 4:**
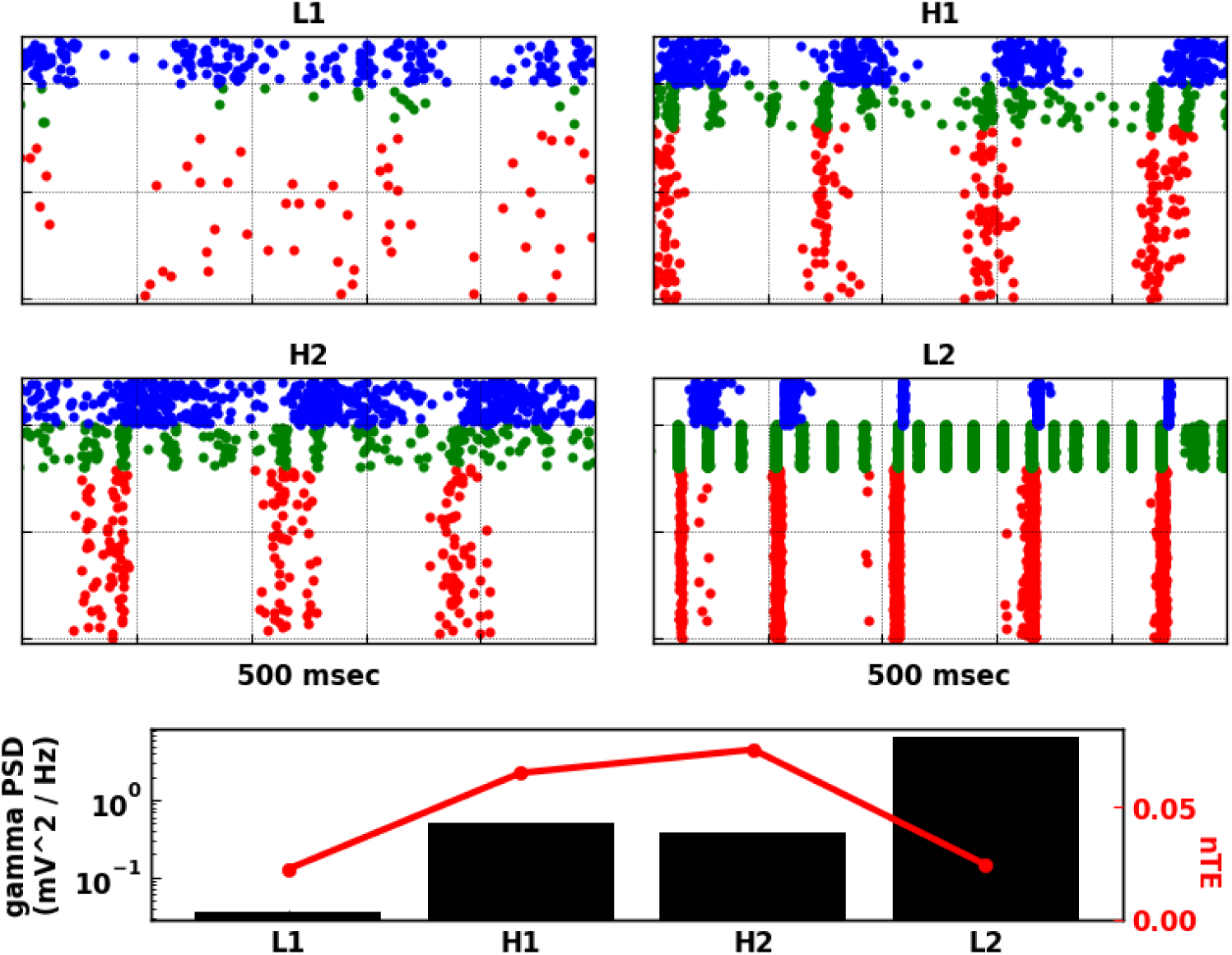
Intermediate excitability with intermediate synchrony allowed for high information transfer across tri-scaling manipulations. Raster plots based on parameters labeled L1, L2 (with reduced information flow), and H1, H2 (with increased information flow) on Fig. 3 for an example simulation. L1 showed low firing; L2 showed high synchrony. H1 and H2 showed similar firing patterns, where both excitability and synchrony were midway in comparison to simulations in L1 and L2. In bottom panel, gamma power reflected the different raster dynamics, with intermediate gamma power for H1 and H2 simulations in comparison to gamma power of simulations L1 and L2. nTE showed an inverted-U pattern with increasing gamma power.

Examination of rasters from points of high information flow-through (H1-H2 in Fig. 3) showed moderate levels of activity (H1-H2 in Fig. 4). Firing here was intermediate between the low-firing of L1 and the strong phase-locking of L2, consistent with the inverted-U pattern. This intermediate synchronization allowed the pyramidal neurons to be excitable enough for the timing of their firing to be appropriately biased by driving input to allow information flow. Gamma power of H1 and H2 in the bottom panel of Fig. 4 was also intermediate between gamma power of L1 and L2. This pattern generalized across all manipulations, where maximum nTE was found at a mid-gamma power range. As shown in Fig. 5, gamma power from the tri-scaling simulations was in the range of 0.01 to 10.8 *mV* ^2^*/*/Hz. However, the simulations with nTE in the top 10^th^ percentile had gamma power at an intermediate range of 0.02 to 2.12 *mV* ^2^/Hz.

**Figure 5:**
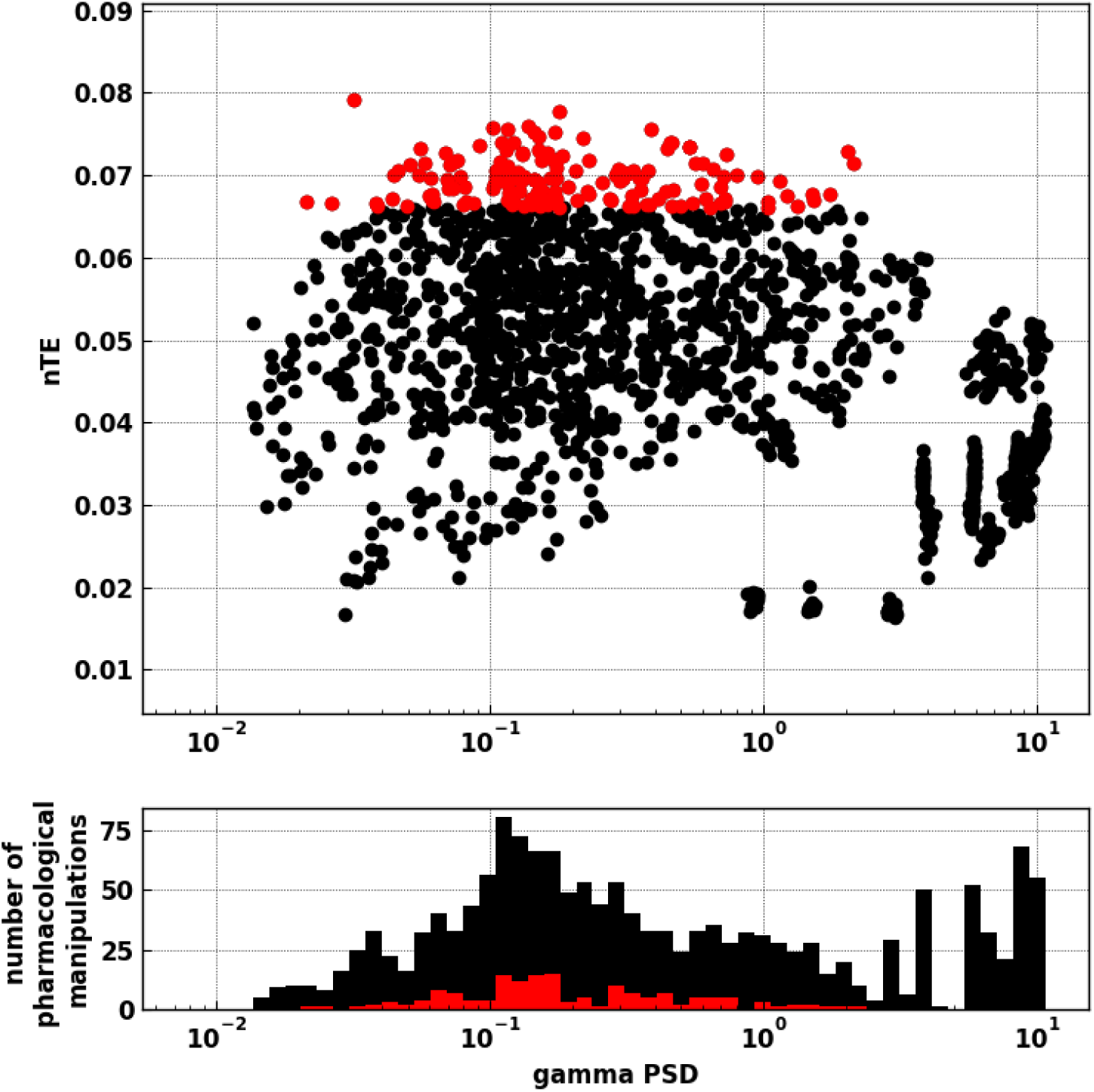
High nTE values were found at intermediate gamma power, suggesting that extremes of gamma power reduced information flow. Top panel shows gamma power versus nTE in simulations from tri-interaction pharmacological manipulations. Each dot represents a simula-tion. The red set represents manipulations with nTE values in the top 10th percentile, which lied in an intermediate gamma power range from 0.02 to 2.12 *mV* ^2^*/*Hz. Bottom panel shows a histogram of the distribution of the data points in the top panel.

The interaction between excitability and synchrony could be seen in the inverted-U shaped relationship of information flow with the firing rate of pyramidal neurons and PV interneurons (Fig. 6). At low firing rate of pyramidal neurons, reflecting low excitability, there was reduced information flow (Fig. 6A, left). As firing rate increased, information flow increased. However, after a certain degree of excitability, information flow started decreasing. This could be explained by the limiting effect of highly synchronized firing on information flow, as reflected in the population synchrony of pyramidal neurons (red markers in Fig. 6C). Population synchrony was estimated using the coefficient of variation from the interspike intervals across all the neurons in a population [88]. Population synchrony ∼ 1 represents interspike intervals that belong to a Poisson distribution, reflecting random population firing driven by the Poisson driving input. In a synchronized population, the upper range of population synchrony approaches 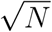, where *N* is the number of neurons firing synchronously at each firing cycle. At highly synchronous firing of pyramidal neurons (red markers in Fig. 6C) and PV interneurons (red markers in Fig. 6D), information flow was reduced. The coloring in A to F of Fig. 6 is for quintiles of pyramidal neuronal population synchrony, to keep track of the simulations with similar manipulations across firing rate and population synchrony.

**Figure 6:**
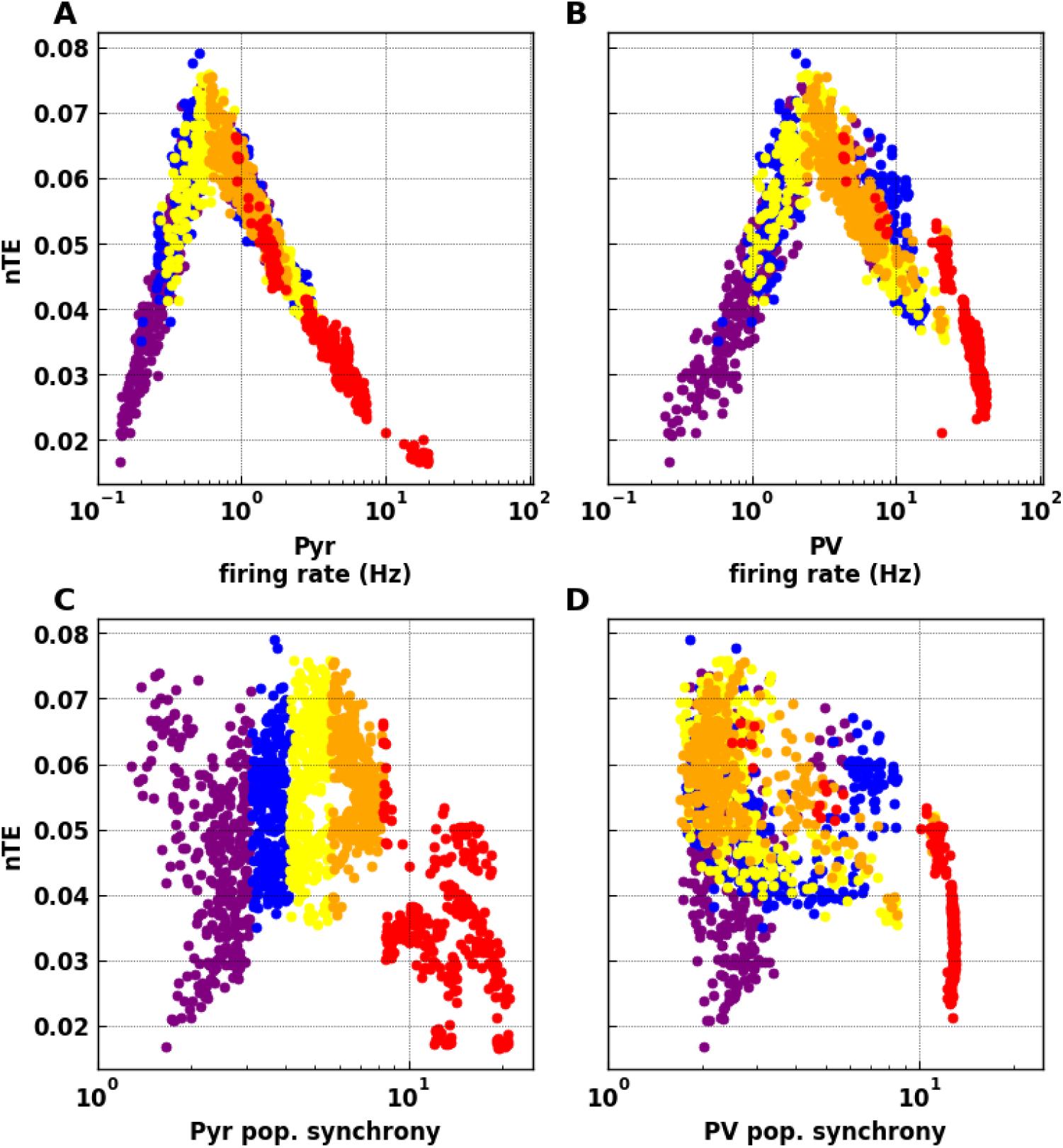
Pyramidal and basket cell population firing rates and synchrony were associated with information flow changes. Variability of population firing measured with population synchrony. Points in the panels, representing individual simulations, are color-coded according to quintiles measured for pyramidal population synchrony. Quintiles show a general consistency across these 4 measures.

## 3 Discussion

Using a computer model of the CA3 microcircuit, we found an inverted-U pattern which related information flow both to potential therapeutic targets schizophrenia: NMDAR, GABA_A_R, and *I*_h_; and to gamma activity. These findings suggest an interesting interaction between pyramidal neuronal excitability and synchrony (Fig. 7). At low excitability, pyramidal neurons were below firing threshold and showed low activity. A driving input was less likely to influence or bias spike timing. Therefore, information flow was low. As excitability increased, more pyramidal neurons were closer to the firing threshold, and the driving input could more strongly bias firing, increasing information flow. At the other extreme of excitability, high pyramidal neuronal activity increased activity of basket interneurons, increasing gamma power via both ING and PING mechanisms. As gamma power got higher, pyramidal cell activity was increasingly locked into the oscillation, reducing the ability of inputs to influence firing times and reducing information flow. These findings provide a mechanistic explanation connecting pyramidal neuronal excitability and activity, with population dynamics and oscillations, in the context of potential therapeutic targets in schizophrenia.

**Figure 7:**
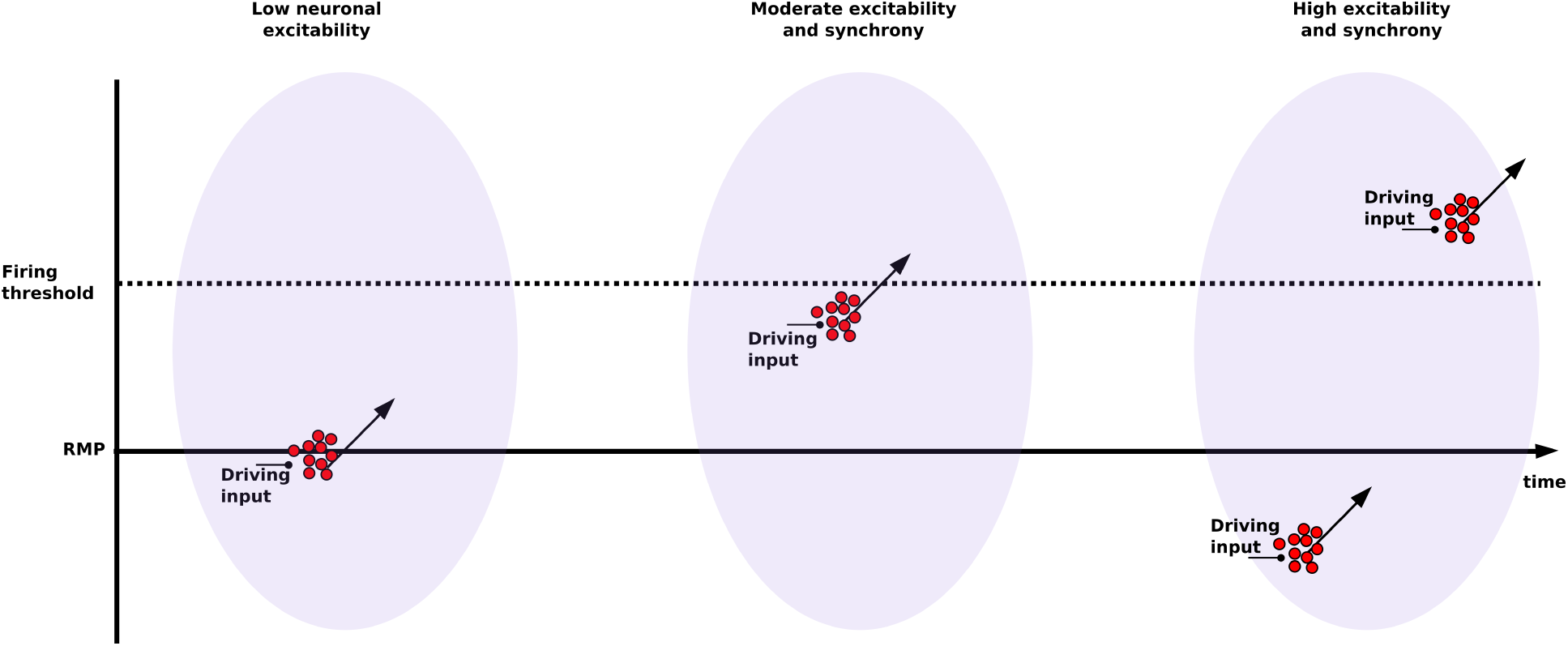
Interaction between pyramidal neuronal excitability and synchrony affected response to driving input, and thus information flow. Driving inputs arriving at a population of pyramidal neurons (red circles) increases excitability (arrows). At *low excitability*, driving input is not enough to reach threshold and trigger firing, reducing information flow from driving input. At *moderate excitability*, pyramidal neurons population is close to firing threshold and so driving input is enough to push cells into firing, At *high excitability*, pyramidal neurons are pushed back-and-forth between synchronized firing with little driving input influence relative to internal drive, and synchronized inhibition with little input influence due to distance from threshold. (RMP: resting membrane potential).

Our study makes the following testable predictions: **1.** Augmenting NMDARs on OLM interneurons, for example using photo-switchable NMDAR GluN2A or GluN2B subunits [5] expressed specifically in the OLM interneurons, will show alterations in gamma activity *in vivo*. **2.** The effect of augmenting NMDARs on information flow will depend on the dynamic state of the network reflected by gamma power. Optogenetic stimulation of channelrhodopsin-2 expressed in the PVs at gamma frequency could be used to manipulate gamma power [9]. Information flow could be measured by nTE using LFPs [33]. **3.** Manipulations of *I*_h_ with photo-activated adenylate cyclase expressed specifically in PVs and PYRs, or by using norepinephrine, will produce our observed changes in gamma and information flow.

### 3.1 Why is information flow important?

Either extreme of information flow will interfere with information processing in the brain. Brain organization is thought to be grossly hierarchical [92]. As a first approximation, we can ignore feedback circuits and consider the processing of information at each level from the prior processing stage [27, 51]. We have previously hypothesized, and demonstrated in a model, that proper information processing at each stage requires a balance between information flow-through and information from the area itself, related to local resonance properties [61]. In the present model, where there is relatively low information flow-through, flow-through maximization would be optimal. Decreased flow-through across the CA3 microcircuit would provide decreased information at CA1, the next step in processing, with activity primarily related to CA3 dynamical state rather than to input from dentate gyrus.

Excessive dependence on internal activity has been proposed as a possible pathophysiological mechanism underlying the development of delusions in schizophrenia. Similar to our conclusions, Tamminga *et al.* [86] suggested that CA3 hyperactivity in schizophrenia patients might indicate that the hypothesized pattern completion function of CA3 is excessive, resulting in faulty pattern completion that is not based on environmental input, resulting in delusions. In the cognitive domain, decreased information flow would impair performance because information conveyed to the next processing step has emerged more from the dynamics of the microcircuit and are less related to task input.

### 3.2 Relation to Other Studies

Coordination between different neural processes (neural coordination), including neuronal firing and oscillations, has been proposed to be important in cognitive coordination required to coordinate two or more frames of reference (*e.g.,* visual and olfactory frames of reference in rodents), whether maintained in separate ensembles simultaneously or provided by alternating ensemble activation at delta or theta frequency [46]. Neural discoordination would then underlie cognitive symptoms (cognitive discoordination) in schizophrenia [47, 68, 91]. Failure to coordinate multiple frames could result in difficulty separating frames reflecting internal processing (expectations, Bayesian priors, imagining, planning) from frames reflecting external realities (stimulus associated). In our model, moderately-synchronized pyramidal neuronal firing was needed to increase information flow.

Impaired excitation/inhibition balance has been proposed to be one of the pathophysiological mechanisms in schizophrenia. In a combined fMRI and computer modeling study, [81] showed impaired working memory deficits suggestive of cortical disinhibition. Impaired PV functioning in schizophrenia suggested the shift of the excitation/inhibition balance towards more excitable microcircuit as well [4, 48, 19]. Our model suggested how more excitable microcircuit can reduce information flow through increased synchrony.

Other groups have studied the relationship between gamma power and information flow outside the context of schizophrenia. Buehlmann et al [7] showed increased transfer entropy with increased gamma power between two sets of excitatory and inhibitory integrate-and-fire neurons. In addition to having a simpler model, they performed a modulation different than what we used here, the 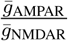 ratio, which may not have permitted them to explore as broad a range of gamma. Akam et al [2] described a model where oscillations provided an additional channel for information flow besides the rate of neuronal firing. They studied the flow of information from four input populations of excitatory and inhibitory neurons into one output population. Switching the dynamics of one input population from irregular to oscillatory firing resulted in additional information flow, consistent with our study.

### 3.3 Limitations of the study

The major limitations of this study are the limitations that are inherent in all modeling studies – we necessarily made choices as to what to include and what to leave out. The things left out include both “known unknowns … and unknown unknowns.” With respect to the unknown unknowns, we have limitations that are comparable, though more severe, than those of other studies – once you move to an animal disease model (*in vivo*) or remove tissue from the organism (*in vitro*) or simply extract parameters (*in silico*), you are eliminating much of the clinical phenomenology. With respect to the known unknowns, we are progressively adding detail and specifics using the best information available, but we continue to have computing and research limitations that reduce the fidelity of the model. In particular, **1.** we omitted interneuron populations other than PV, OLM cells; **2.** we omitted dopaminergic and serotonergic receptors, the targets of most current antipsychotic medications; **3.** we modeled PV and OLM as single compartments without 3D details; **4.** there is inadequate information regarding the distribution of voltage- and calcium-sensitive ion channels in PYR dendrites.

### 3.4 Clinical relevance

Our multiscale modeling study suggests that network synchronization and pyramidal neuronal excitability are potential dynamical targets for the treatment of cognitive symptoms and information processing deficits in schizophrenia. We showed how manipulating multiple molecular elements in a multi-target pharmacotherapy approach explains some of the inverted-U shaped phenomena seen in schizophrenia [43]. Multi-target pharmacotherapy is necessarily already used in schizophrenia treatment. Clozapine, often used after the failure of multiple other antipsychotic medications [67], is considered a “dirty” medication that targets a wide array of receptors [72]. A multi-target pharmacotherapy approach is also supported by the evidence that multiple genes and proteins have been identified in the pathophysiology of schizophrenia (*e.g.,* see [95]).

A number of FDA approved pharmacological agents act on the molecular targets we investigated here, and so could be explored for multi-target drug therapy complementing NMDAR augmentation in the treatment of cognitive symptoms in schizophrenia. GABA_A_R modulators include benzodiazepines, which are not subunit-specific, as well as subunit-specific agents, *e.g.,* zolpidem [76]. But until now, GABA_A_R modulators are not neuronal subpopulation specific. In regard to *I*_h_, two medications, lamotrigine and gabapentin, used for various neurological and psychiatric disorders, upregulate HCN1 levels [70].

Our study points to a possible explanation of an non-intuitive relationship between gamma power and symptoms of schizophrenia. In schizophrenia and pharmacological models of schizophrenia, gamma power has been found to be decreased [90, 91, 74] or increased [30, 29, 23], depending on the study. Our results suggest a way to resolve the paradox: both extremes of gamma power interfere with information processing. Psychotic symptoms, such as delusions of control and hallucinations, as well as cognitive symptoms, have been conceptualized as being due to “dysconnection” syndromes [83], where the communication between different brain regions is disrupted. We would instead suggest that the complexity and variability of psychotic manifestations might instead be due to re-connection or rewiring syndromes, where areas of decreased oscillatory strength will be disconnected and areas of increased oscillatory strength hyper-connected but with little variability and responsiveness to outside inputs (stereotypic thoughts and behaviors). This rewiring would be due to areas being pushed out of an essential central functional regime of gamma power. This further suggests the importance of careful titration when using schizophrenia medication; it would be easy to overshoot a target gamma-power zone.

## 4 Methods

The full model is available on modelDB#258738 as http://modeldb.yale.edu/258738. It is largely based on [64, 66]; modelDB#139421. The model was implemented in NEURON 7.4 [10] running in parallel on eight 2.67 GHz Intel Xeon quad-core CPUs [28]. Result robustness was tested by using multiple random wirings and random stimulation input patterns for each parameter set, resulting in around 1900 simulations.

The model consisted of 800 pyramidal (PYR) neurons, 200 oriens-lacunosum moleculare (OLM) interneurons, and 200 parvalbumin-positive (PV) fast-spiking basket interneurons (Fig. 8), all modeled using multiple channels defined using Hodgkin-Huxley parameterizations. PYRs had five compartments comprising soma, apical dendrite, and basal dendrite. OLM and PV cells were single compartment. All neurons contained leak, fast sodium, and delayed rectifier, *I*_h_. PYRs also had A-current, and had increasing *I*_h_ conductance up apical dendrite [39]. OLM added Ca^2+^ -gated K^+^ current and high-threshold Ca^2+^ current with intracellular calcium dynamics.

**Figure 8:**
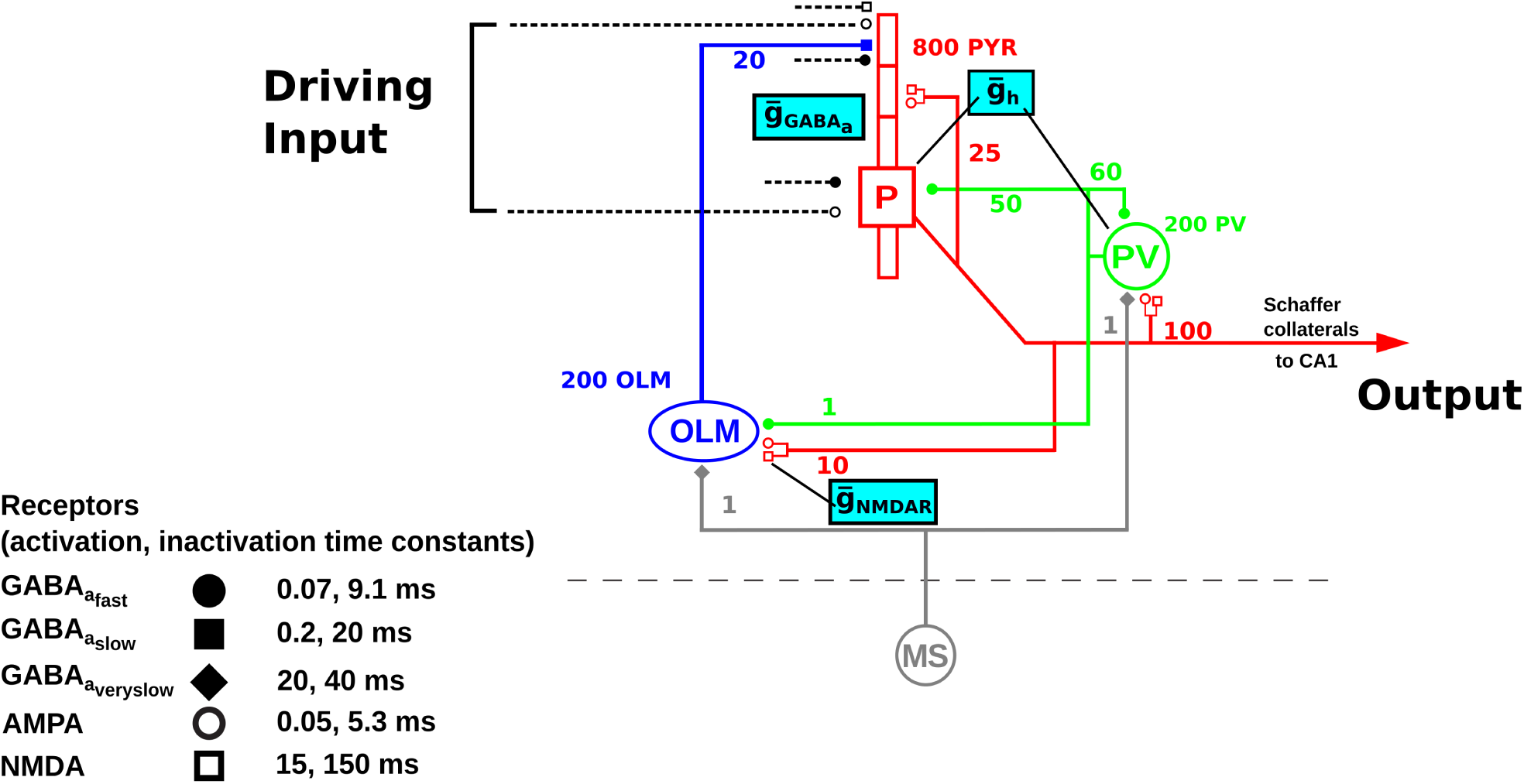
Schematic diagram for the *in silico* CA3 network model showing targets for multidrug target manipulations. Population of pyramidal neurons (n = 800) is represented by the red cell, PV basket interneurons (n = 200) by the green cell, and OLM interneurons (n = 200) by the blue cell. Dotted lines represent the random input driving the pyramidal neuron population (driving inputs). Output represents the output of pyramidal neurons. MS is medial septum, providing inhibitory input onto the PV basket and OLM interneurons. Numbers on connections are convergence ratios for connectivity between different populations. Cyan boxes show the molecular targets that are being investigated in this study and their locations: 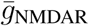 on OLM interneurons, 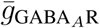 on pyramidal neurons, and 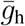 on pyramidal neurons and PV basket interneurons. Activation and inactivation time constants for each of the receptor types are shown.

PYRs projected both AMPARs and NMDARs on all cell types, with mid-apical projections onto other PYRs. PVs and OLMs projected to GABA_A_R: PV → PYR soma; OLM → PYR distal apical. OLM, PV received GABA_A_R input from medial septum (MS) at 6.7 Hz (theta). Synapses were modeled as double exponential mechanisms with parameters from [89] (Table 1). Background activity was simulated low amplitude Poisson input to all populations [17] 2. Five seconds of simulation time were used for analysis, after discarding the first two seconds to allow network dynamics to stabilize. Local field potential (LFP) was calculated as voltage difference between apical and basal dendrites of all PYRs.

**Table 1:** Synaptic parameters for neuronal connectivity

| Presynaptic | Postsynaptic | Receptor | $\tau_1$ (ms) | $\tau_2$ (ms) | Conductance (nS) | Convergence |
| --- | --- | --- | --- | --- | --- | --- |
| Pyramidal | Pyramidal | AMPA | 0.05 | 5.3 | 0.02 | 25 |
| Pyramidal | Pyramidal | NMDA | 15 | 150 | 0.004 | 25 |
| Pyramidal | Basket | AMPA | 0.05 | 5.3 | 0.36 | 100 |
| Pyramidal | Basket | NMDA | 15 | 150 | 1.38 | 100 |
| Pyramidal | OLM | AMPA | 0.05 | 5.3 | 0.36 | 10 |
| Pyramidal | OLM | NMDA | 15 | 150 | 0.7 | 10 |
| Basket | Pyramidal | GABA<sub>A</sub> | 0.07 | 9.1 | 0.72 | 50 |
| Basket | Basket | GABA<sub>A</sub> | 0.07 | 9.1 | 4.5 | 60 |
| Basket | OLM | GABA<sub>A</sub> | 0.07 | 9.1 | 0.023 | 1 |
| OLM | Pyramidal | GABA<sub>A</sub> | 0.2 | 20 | 72 | 20 |

**Table 2:** Synaptic parameters for modeling background activity

| Cell | Section | Synapse | $\tau_1$ (ms) | $\tau_2$ (ms) | Conductance (nS) |
| --- | --- | --- | --- | --- | --- |
| Pyramidal | Soma | AMPA | 0.05 | 5.3 | 0.05 |
| Pyramidal | Soma | GABA<sub>A</sub> | 0.07 | 9.1 | 0.012 |
| Pyramidal | Dend | AMPA | 0.05 | 5.3 | 0.05 |
| Pyramidal | Dend | NMDA | 15 | 150 | 6.5 |
| Pyramidal | Dend | GABA<sub>A</sub> | 0.07 | 9.1 | 0.012 |
| Basket | Soma | AMPA | 0.05 | 5.3 | 0.02 |
| Basket | Soma | GABA<sub>A</sub> | 0.07 | 9.1 | 0.2 |
| OLM | Soma | AMPA | 0.05 | 5.3 | 0.0625 |
| OLM | Soma | GABA<sub>A</sub> | 0.07 | 9.1 | 0.2 |

Target locations for NMDAR and *I*_h_ pathological manipulations were based on our prior work: NMDAR on OLM: modelDB#139421 [66, 45]; *I*_h_ on PYR, PV modelDB#151282 [64]. We manipulated GABA_A_Rs on PYRs based on localization evidence from mRNA study in schizophrenia [25].

LFP power spectrum density (PSD) was calculated using Welch method (Python Scipy signal module) [32] after removing DC component. Gamma power was measured as a 25-50 Hz band. Normalized transfer entropy (nTE) was used to measure information flow with 15 ms binning [21, 65]; see ModelDB#136095 and Fig. 9. Firing synchrony was measured using population coefficient of variance (popCV) [88]: 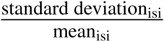 for interspike intervals (isi) for all PYR spikes [88].

**Figure 9:**
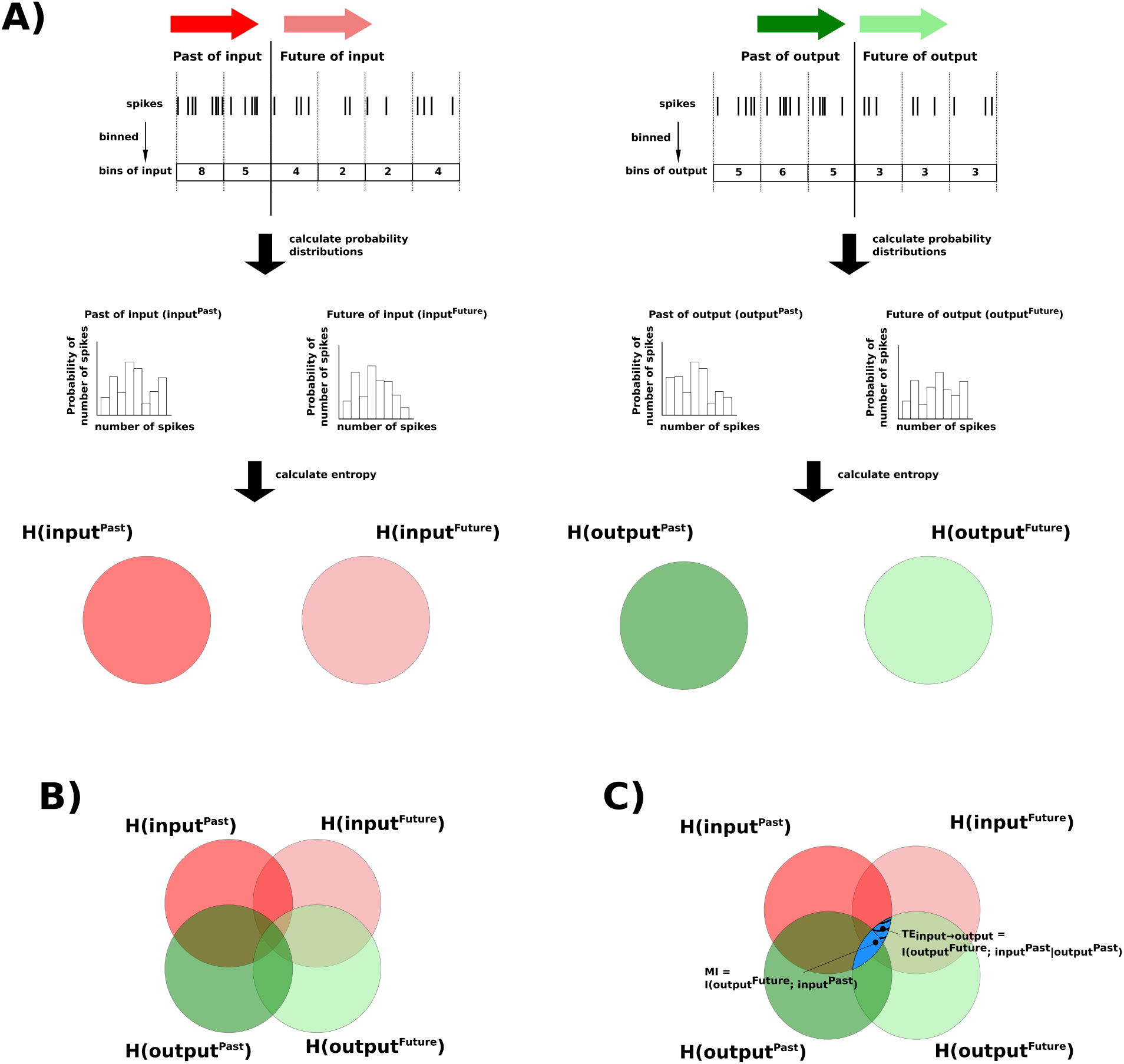
Schematic diagram for steps to calculate transfer entropy (TE) from input to output spikes of a single pyramidal neuron. **A)** Input signal is represented by red arrows (darker red for Past, lighter red for Future). Output signal is represented by green arrows (darker green for Past, lighter green for Future). Spikes in both input and output signals were binned, and the spike counts were used to generate probability distributions to calculate entropy. **B)** The entropies of the signals are overlapped, represented by areas of overlap of the sets in the Venn diagram. The overlap of the entropies is partly because of the flow of information between the different signals, and partly because of chance. **C)** The blue region, representing the overlap between *H*(input^Past^) and *H*(output^Future^), is the mutual information (MI) between the Past of the input signal and the Future of the output signal (*I*(*H*(input^Past^); *H*(output^Future^))). The striped portion of the blue region represents TE from the Past of the input signal to the Future of the output signal. TE was calculated as the MI between *H*(input^Past^) and *H*(output^Future^), given the Past of the output signal (*I*(*H*(input^Past^); *H*(output^Future^)*|H*(output^Past^))).

## 5 Acknowledgments

Supported by: The department of Veterans Affairs, Veterans Health Administration, VISN 1 Career Development Award, and VA Special Psychopharmacology Research Fellowship Program, and New York State Tuition scholarship for graduate students to Mohamed A. Sherif; IMAG (U01EB017695) and NIMH (R01MH086638) to William W. Lytton; and NIDCD (R01DC012947-06A1) and Army Research Office Grant (W911NF-19-1-0402) to Samuel A Neymotin. The authors declare they have no financial conflicts of interst in relation to the work presented. We would like to thank Larry Eberle (SUNY Downstate) for Neurosim lab support; Tom Morse and Ted Carnevale (Yale) for ModelDB support; and the Shepherd lab (Yale) for helpful comments. The views and conclusions contained in this document are those of the authors and should not be interpreted as representing the official policies, either expressed or implied, of the Army Research Office or the U.S. Government. The U.S. Government is authorized to reproduce and distribute reprints for Government purposes notwithstanding any copyright notation herein

## Notes

#### Summary of Updates

Changed order to have methods at the end - updated figure of rasters and gamma power.

